# Genetic analysis of *TP53* gene mutations in exon 4 and exon 8 among esophageal cancer patients in Sudan

**DOI:** 10.1101/572214

**Authors:** Sulafa Mohamed Eltaher, Abeer Babiker Idris, A. H Mahmoud, Mawadah Yousif Mohamed Yousif, Nouh Saad Mohamed, Muzamil M. Abdel Hamid, Kamal Elzaki Elsiddig, Mohamed A. Hassan, Galal Mohammed Yousif

**Affiliations:** The Academy of Health Sciences, The Republic of Sudan Federal Ministry of Health, Khartoum, Sudan; Department of medical microbiology, Faculty of Medical Laboratory Sciences, University of Khartoum, Khartoum, Sudan; Applied Bioinformatics Center, Africa City of Technology, Khartoum, Sudan; Department of Parasitology and Medical Entomology, Institute of Endemic Diseases, University of Khartoum, Khartoum, Sudan; Department of Molecular Biology, National University Biomedical Research Institute, National University, Khartoum, Sudan; Department of Parasitology and Medical Entomology, Faculty of Medicine, Sinnar University, Sinnar, Sudan; Department of Parasitology and Medical Entomology, Nile College, Khartoum, Sudan; Department of Surgery, Faculty of Medicine, University of Khartoum, Khartoum, Sudan; Department of Bioinformatics, DETAGEN Genetic Diagnostics Center, Kayseri, Turkey; Faculty of Pharmacy, Alrebat University, Khartoum, Sudan

**Keywords:** Esophageal carcinoma, Small cell carcinoma, SNPs, functional analysis, in silico tools

## Abstract

**Background:** Esophageal carcinoma (EC) represents the 1^st^ rank among all gastrointestinal cancers in Sudan. Despite little publications, there is a deep absence of literature about the molecular pathogenesis of EC considering TP53 gene from Sudanese population.

**Aims:** In this study, we performed the expression analysis on p53 protein level by immunohistochemical staining and examined its overexpression with p53 mutations in exons 4 and 8 among esophageal cancer patients in Sudan.

**Material and Methods:** Fixed tissue with 10% buffered formalin was stained by Hematoxlin and Eosin (H&E), Alcian blue-Periodic Acid Schiff (PAS) and Immunohistochemistry stain. PCR-RFLP was used to study the frequencies of p53 codon 72 R/P polymorphism. Conventional PCR and sanger sequencing were applied for exon 4 and exon 8. Then detection and functional analysis of SNPs and mutations were performed using various in bioinformatics tools.

**Result:** Nuclear accumulations for p53 protein was detected in all of the esophageal carcinomas examined while no accumulations were observed in normal control sections. Four patients with immune-positive for p53 showed no mutations in p53 gene (exon4 and exon8). The incidence of the homozygous mutant variant Pro/Pro was higher in esophageal cancerous patients comparing to healthy control subject 20(71. 4%) vs. 1(10%), respectively (p=0.0026). In exon 4, no mutation was detected other than NG_017013.2:g. 16397C>G. While in exon 8, g.18783-18784AG>TT, g.18803A>C, g.18860A>C, g.18845A>T and g.18863_ 18864 InsT were observed.

**Conclusion:** we found a significant association between the overexpression of TP53 protein and mutation in exon 4 and 8. A silent mutation P301P was detected in all of examined cases. Two patients who diagnosed with small cell sarcoma have shared the same mutations in exon8. Further studies with large sample size are required to demonstrate the usefulness of these mutations in the screening of EC especially SCCE.

## 1. Introduction

Esophageal carcinoma (EC) is the 8^th^ most common diagnosed cancer worldwide and the 6^th^ leading cause of cancer related mortality.(1–3) EC primarily happens in one of two forms: esophageal squamous cell carcinoma (ESCC), which is more prevalent in developing countries, arising from the stratified squamous epithelial lining of the organ; and esophageal adenocarcinoma (EAC) that is a distal esophageal cancer and arising from a metaplastic transformation of the native esophageal squamous epithelium into columnar epithelium due to known risk factors, i.e., obesity, smoking, gastroesophageal reflux and Barrett’s esophagus (BE).(4–6) ESCC is mainly associated with multiple factors such as smoking, alcohol consumption, hot tea drinking, red meat consumption, poor oral health, low intake of fresh fruit and vegetables, and low socioeconomic status. Recently, there has been an increase in the incidence of EAC, especially in developed countries.(4, 7, 8) This arising could be due to multiple factors, such as environmental, together with existing genetic susceptibility factors.(9) Sarcomas and small cell carcinomas usually constitute less than 1-2% of all esophageal cancers.(10) In a logical attempt to understand the remarkable diversity of neoplastic diseases, Hanahan and Weinberg have proposed eight hallmarks of cancer, more over they added two enabling characteristics that make the acquisition of these hallmarks possible: genome instability and mutation, and tumor-promoting inflammation.(11, 12) Knowing about these concept may lead to develop a new approaches to treat human cancers. Many of studies have suggested that the polymorphisms in functionally critical genes may be involved in esophageal carcinoma (13). The most important genes are those which act as anti-oncogenesis. Loss of function for these genes may be even more important than proto-oncogene/oncogene activation for the process of esophageal oncogenesis.(14) The most important tumor suppressor gene that are reported widely in association with different types of cancer is *p53* gene. (15)

*TP53* gene (ID: 7157, MIM: 191170) is often referred to as “the guardian” of the human genome. It mapped on 17p13 and composed by 11 exons (∼20 KB) encoding a nuclear p53 protein of 393 amino acids.(15–17) This regulatory protein controls the expression of hundreds of genes and noncoding RNAs, as well as the RNA processing complexes activity. Also, p53 involves in the checkpoint at the G1/S boundary of cell growth cycle and preventing the multiplication of damaged cells.(16, 18–20) However, The p53 protein has others biological functions, for example, senescence, DNA metabolism, angiogenesis, cellular differentiation, and the immune response.(15) Single nucleotide polymorphisms (SNPs) of TP53 gene are expected to cause measurable perturbation on p53 function. These genetics variants in TP53 implicated in the development of cancer because they are supposed to influence cell cycle progression, apoptosis, and DNA repair.(15) At least 85 SNPs are reported on TP53. The common missense (non-synonymous) polymorphism occurs at codon 72 of exon 4 in the transactivation proline-rich domain of the protein where either CCC encodes proline or CGC encodes arginine (TP53 Arg72Pro, rs 1042522).(21) Some studies have investigated the association of Arg72Pro polymorphism with different kinds of cancers such as esophageal (22), gastric (15), colorectal (23), lung (24), cervical and breast cancer. (25)

In Sudan, EC is a growing problem. In earlier study, conducted in Khartoum during the period 1965 to 1974, reported that the incidence of EC was 1.4% of all malignant tumors. (26) In contrast, a study conducted in Gezira province, in central Sudan, during the period from January 2005 to December 2006 revealed that 9.6% of patients referred for endoscopy proved to have esophageal cancer. (27) Now, EC represents the 1st rank among all gastrointestinal (GI) cancers in Sudan. (10, 28) Unfortunately, there is a deep absence of literature talking about the molecular pathogenesis of EC considering p53 gene in Sudan, despite little publications. Therefore, here we investigated the association between the overexpression of TP53 protein and mutation in exon 4 and exon 8. Then studied their roles in tumorigenesis using in silico tools. To the best of our knowledge, in this study, for the first time in Sudan, we analyzed the genetic alterations in exon 4 and exon 8 of patients with small cell sarcoma of esophagus (14.29% of our patients).

## 2. Materials and methods

### 2.2 Study population

This study included 24 primary esophagus carcinoma patients. All recruited from department of endoscopic. Tumor types and stages were determined by experienced pathologists. Blood samples of 20-age and gender –matched cases with no signs of any malignancy were collected as controls. The mean age of both patients and control groups was 50 years old and 15 patients and14 controls were >50 years old. Data on all esophagus carcinoma were obtained from personal interviews with patients and or co-patients, medical records and pathology reports. The data collected included gender, age, dwelling, tumor location, symptoms and risk factor exposure. The demographic characteristics of the cases and controls; and clinicopathological characteristics of cases are summarized in Table 1 and Table 2.

**Table 1:**
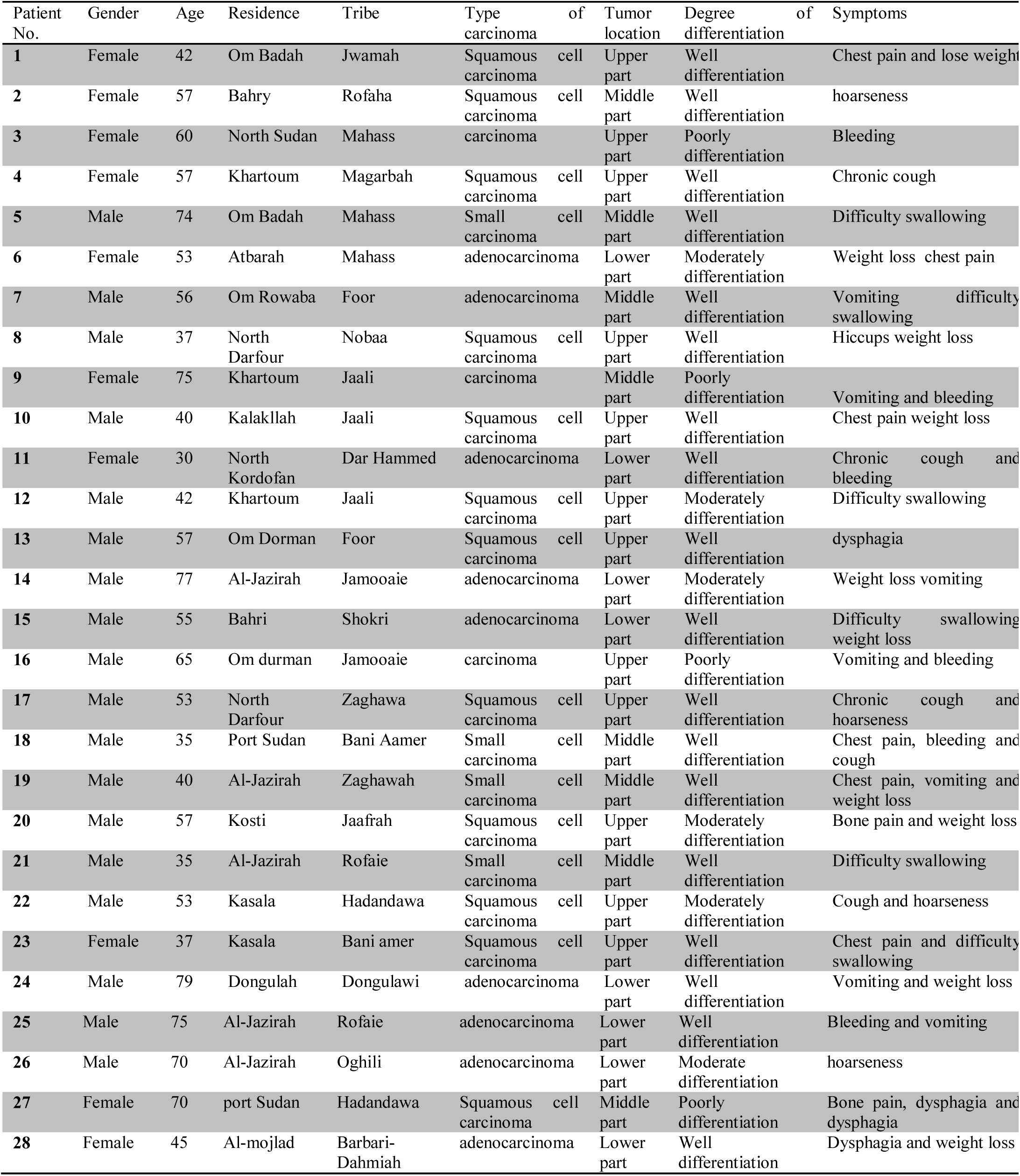
Demographic and clinicopathological characteristics of esophageal cancer patients

**Table 2.** Demographic characteristics of healthy controls

| Control No. | Gender | Age | Residence | Tribe |
| --- | --- | --- | --- | --- |
| 1 | Male | 35 | North Darfor | Jamiaie |
| 2 | Female | 50 | Bahri | Mahasia |
| 3 | Female | 48 | Khartoum | Mahesia |
| 4 | Female | 80 | North State | Mahesia |
| 5 | Male | 32 | Senar | Jamiaie |
| 6 | Female | 45 | North Kordufan | Jamiaie |
| 7 | Male | 78 | Khartoum | Ababdah |
| 8 | Female | 60 | Khartoum | Magarbah |
| 9 | Male | 60 | Sinar | Hamar |
| 10 | Male | 47 | Om durman | Jafrah |
| 11 | Female | 47 | Al-Jazirah | Hasaniah |
| 12 | Female | 28 | Sinar | Jafrah |
| 13 | Female | 42 | Bahri | Jaali |
| 14 | Male | 40 | Khartoum | Jaali |
| 15 | Male | 62 | Khartoum | Jaali |
| 16 | Female | 30 | Sinar | Daishah |
| 17 | Male | 55 | Shandi | Jaali |
| 18 | Female | 48 | North State | Shigiah |
| 19 | Female | 52 | Shandi | Jaali |
| 20 | Female | 46 | Bahri | Robatab |

All patients and/or co-patient were informed about the study and their consent to participate in this study was obtained. This study was approved by the Ethics Committee of tropical and Sudan academic; and informed consent was obtained from the participants.

### 2.3 Histological analysis

Fixed tissue with 10% buffered formalin was stained by Hematoxlin and Eosin (H&E), carcinoma diagnosis was confirmed by pathologist who looked for the degree of histological differentiation; well, moderate, poor, or undifferentiated tumors. The anatomic subsides were categorized as first third esophagus, the middle third, Lower third and junctional area.

Also, Alcian blue-Periodic Acid Schiff (PAS) stain was used to differentiate between neutral and acetic mucosubstances. We used an internal control of normal tissue mucin control.

### 2.4 Immunohistochemistry

TP53 Immunohistochemistry was performed with the mouse monoclonal antibody Do-7 (Dako, Glostrup, Denmark), according to standard protocols.

### 2.5 Molecular Genetics analysis

#### DNA extraction

DNA was extracted from fresh tissue by using Guanidine chloride method as previously described by Coleman *et al*.(29) The Concentration of DNA was determined Spectrophotometrically.

#### Polymerase chain reaction (PCR)

Extracted DNA was amplified for the TP53 gene. The primers for exon 4 were: forward, 5′TCCCCCTTGCCGTCCCAA3′; reverse, 5′CGTGCAAGTCACAGACTT3′ and for exon 8 were Forward: 5’GGGAGTAGATGGAGCCTGGT3’; reverse: 5’GCTTCTTGTCCTGCTTGCTT3’.(30) Exon 4and 8 of the p53 gene were amplified separately by incubating on cycler for 10min at 94c for initial denaturation followed by 35c cycles at 9ac for 30s, 55c for 30s and 72c for1min. the final extension step was 72c for 7min. After that prepare gel run (Agarose 1.5 gm) run the PCR product on agarose gel visualize the PCR product under UV light.

#### Restriction fragment length polymorphism (PCR-RFLP) analysis

For genotyping of p53 for the codon 72 polymorphism, added 10µl of enzyme *BstUI* and incubated overnight at 37°C. Then the digested product was separated on 3% agarose gel with ethidium bromide and photographed with an Ultra Violet Product Image Store system.(30)

#### DNA sequencing

Out of 48 PCR products, 13 patients and 10 controls PCR products were sent for Sanger dideoxy sequencing, including both forward and reverse nucleotide sequencing, which performed by Macrogen Company (Seoul, South Korea).

#### Sequence analysis

Sequence analysis was done by using Finch TV program version 1.4.0(31). The two chromatograms for each individual, (forward and reverse), were visualized and checked for quality; and The Basic Local Alignment Search Tool (BLAST; https://blast.ncbi.nlm.nih.gov/Blast.cgi) was used to assess nucleotide and protein sequence similarities (32).

#### SNPs detection

Gene Screen software(33) was used for searching about mutations and SNPs in all ABI trace files when compared with a reference sequence (TP53 NCBI Reference Sequence: NG_017013)(34) and calculating alleles frequencies. Then tested sequences with the reference sequence and high similarity sequences (U94788) and (X54156), were obtained from NCBI database and added as control sequences, were aligned to confirm the presence of nucleotide changes by using BioEdit software(35). Finally, by using online ExPASy translate tool(36), all tested sequences were translated to amino acid sequences and compared all together with reference sequence (ID:P04637) using BioEdit software.

#### Functional analysis of SNPs

Selected SNP was predicted functionally by using four online softwares: (1) Sorting intolerant from tolerant (SIFT) software(37) which predicted whether the SNP affects protein function based on sequence homology and the physical properties of amino acids. (2) Polymorphism Phenotyping v2 (PolyPhen-2) software(38) that predicted possible impact of the SNP on the structure and function of a human protein using straightforward physical and comparative considerations. (3) Project hope software(39) which gave analyze the effect of the SNP on the protein structure. (4) I-Mutant software(40) which is used to assess the stability of the SNP-involved protein. (5) Predictor of human Deleterious Single Nucleotide Polymorphisms (PhD-SNP)(41) is predicted the relatedness of SNP or mutation to a disease based on a single SVM trained and tested on protein sequence and profile information.

#### Modeling (3D structure)

The protein sequence of *TP53* gene (ID:P04637) was sent to RaptorX Property, a web server, (http://raptorx2.uchicago.edu/StructurePropertyPred/predict/)(42), to predict structure properties of these protein sequences. Then the 3D structures were visualized by using UCSF Chimera (version 1.8) that is currently available within the Chimera package and available from the chimera web site (http://www.cgl.ucsf.edu/cimera)(43).

#### Statistical analysis

Results of *p53* codon 72 SNP among cancerous patients and controls were analyzed using X^2^ test. P<0.05 was considered statistically significant. Statistical analyses in this study were performed using GraphPad Prism (version 5.0).

## 3. Result

### 3.1 Histopathological result

The microscopic morphology of most slides (Hematoxylin and Eosin) at squamous cell carcinoma were moderately to well differentiate showed that pleomorphic variation in size and shape both in cells and nuclei, abnormal nuclear morphology hyperchromatic contain an abundance of chromatin. Nuclear shape was variable with chromatin clumped and large nucleoli. Large numbers of mitoses with higher proliferative activity appeared in abnormal locations within epithelium cells and Stroma. Well and Moderately differentiated Squamous cell carcinoma showed some bridges and nests of keratin pearls also invasion into the submucosa, poorly differentiated cell carcinoma revealed spreading (of malignant cells seem like spindle cell)at the layers without differentiation highly bizarre mitotic, so special stain PAS (periodic acid shiffs) done to characterization carcinoma from sarcoma.

While in adenocarcinoma of esophagus, most tumors are mucin-producing glandular tumors showing intestinal –type features, in the keeping with the morphology of preexisting metaplastic mucosa. The diffuse type showed poorly differentiated adenocarcinoma with little or no discernible gland formation, tumor cell forming a diffuse sheet infiltrating between bundle of smooth muscle. (Figure1-3)

**Figure1.**
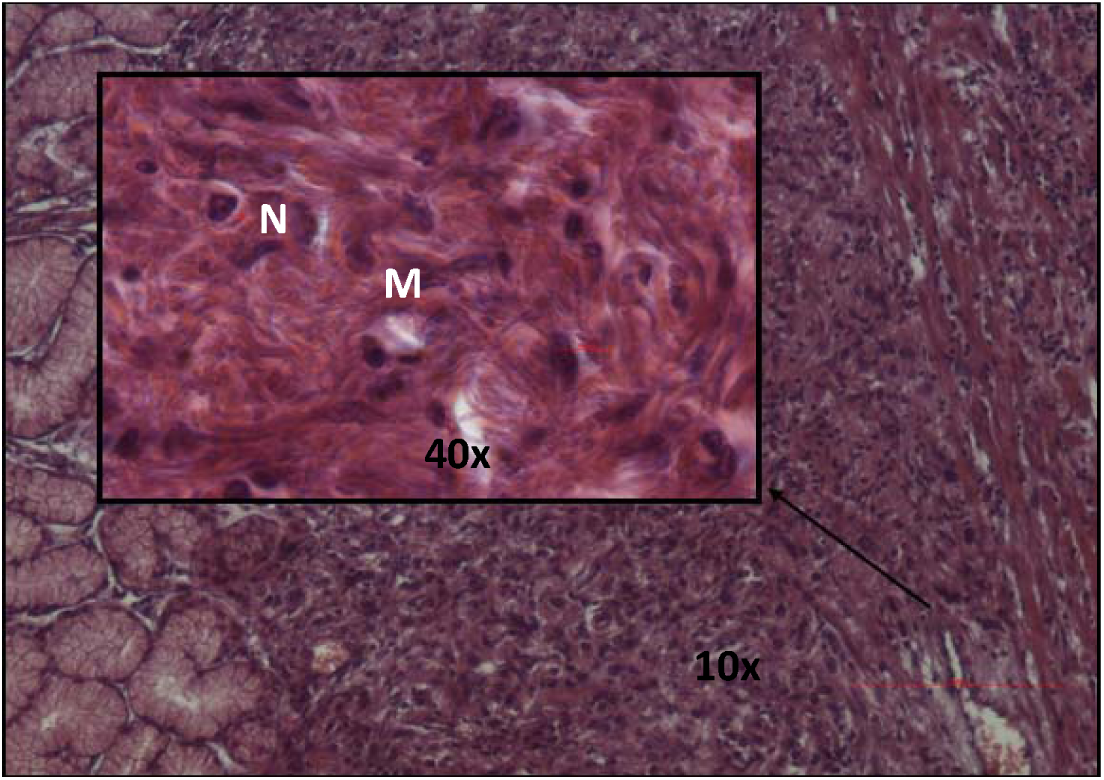
Shows H&E for poorly differentiated carcinoma under the glands by light microscopy. N: indicates nuclei variable in shape and size than in normal (pleomorphism). M: shows high mitotic activity with abnormal mitotic figures (arrows) in an ESCC.

**Figure2.**
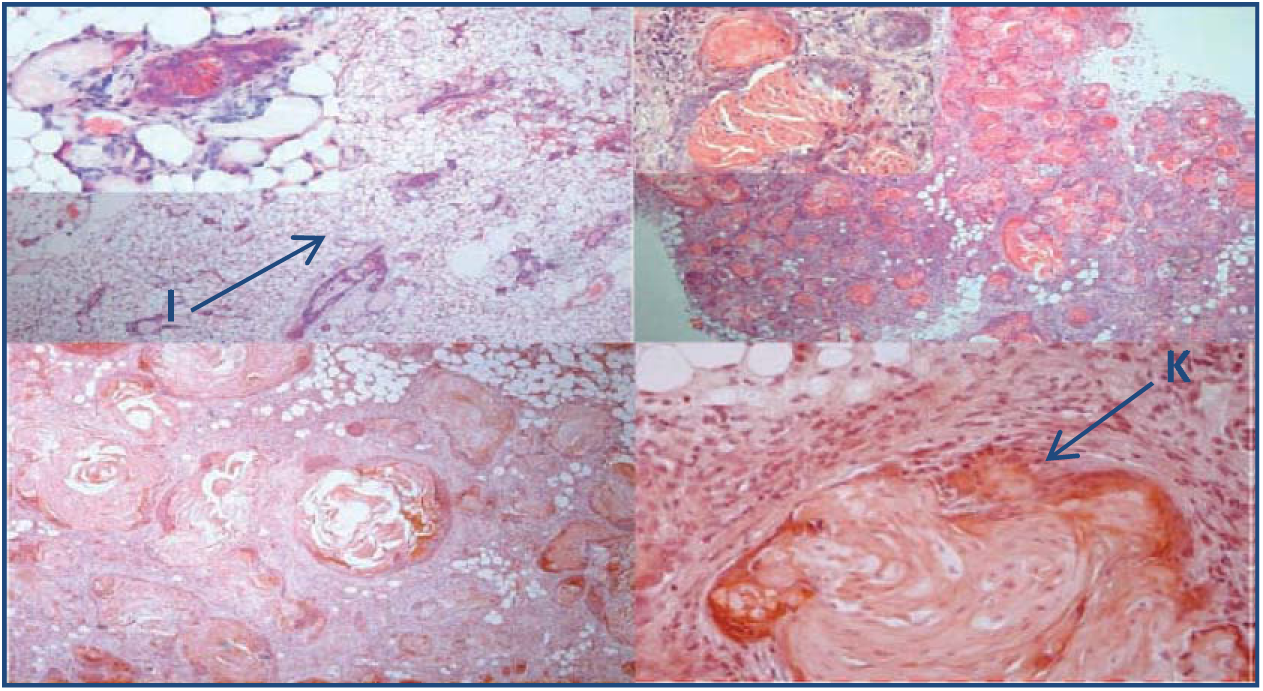
Illustrates well differentiation SCC. showing a hyper proliferative epidermal cyst containing ghost cells and keratinized structures(K), accompanied by an acute stromal inflammatory reaction (I) and epithelial structures.

**Figure3.**
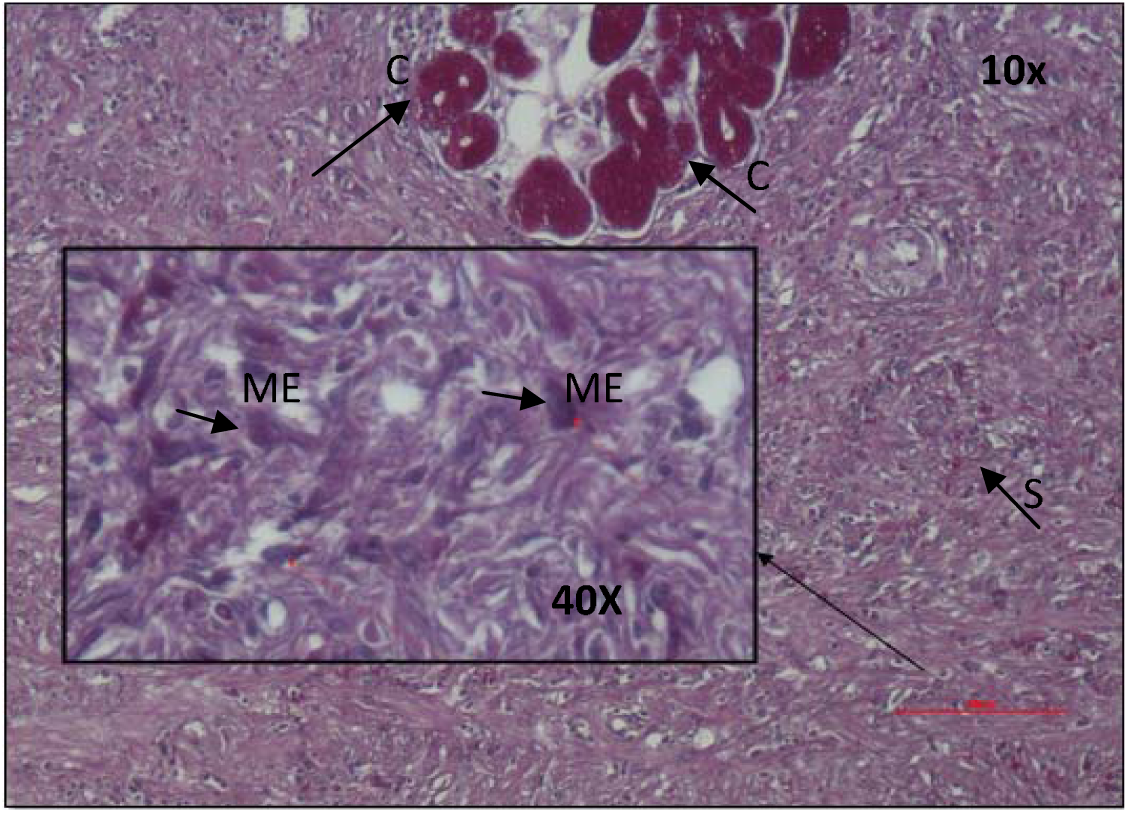
Shows Compound Alcian blue and PAS special stains. Malignant epithelial contains neutral mucin (ME) which seen in malignant nucleolus and stained blue by Alcian blue stain; spreading in stroma(S) contains other type of epithelial cell containing acetic. Mucosubstances (C) are stained magenta color by PAS stain.

### 3.2 Immunohistochemical P53 overexpression

Nuclear accumulations for p53 protein was detected in all of the esophageal carcinomas examined, as illustrated in Figure4. While no accumulations were observed in normal control sections. Four patients (patient1, patient3, patient6 and patient24) with immune-positive for p53 showed no mutations in p53 gene (exon4 and exon8).

**Figure 4:**
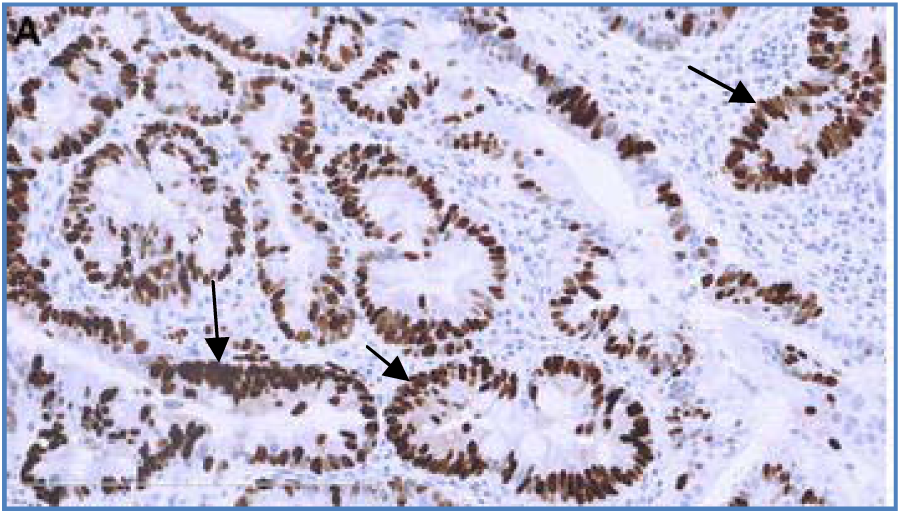
Shows immunohistochemistry expression for p53. No accumulation in mucosa but nuclear accumulation of p53 protein. (black arrows point to the nuclear accumulation in a dark brown color)

### 3.2 PCR-RFLP

*PCR-RFLP* was used to investigate the *p53* codon 72 SNP (dbSNP:rs1042522). The incidence of the homozygous mutant variant Pro/Pro was higher in esophageal cancerous patients comparing to healthy control subject 20(71. 4%) vs. 1(10%), respectively (p=0.0026). (Table3) (Figure 6b)

**Table 3:**
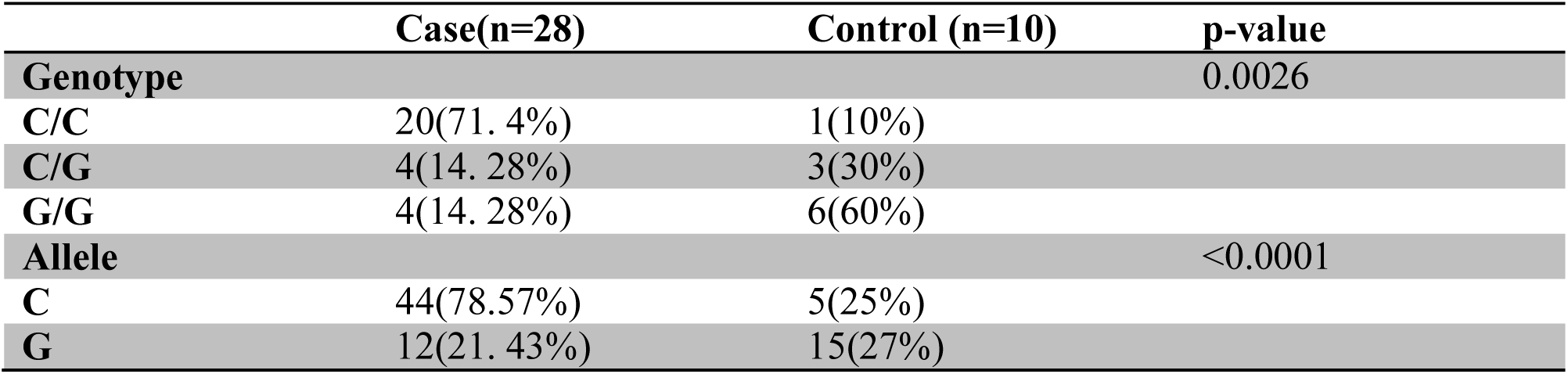
Genotype and allele frequency of the p53 codon 72 SNP in esophageal cancerous patients and controls

|  | Case(n=28) | Control (n=10) | p-value |
| --- | --- | --- | --- |
| **Genotype** |  |  | 0.0026 |
| **C/C** | 20(71. 4%) | 1(10%) |  |
| **C/G** | 4(14. 28%) | 3(30%) |  |
| **G/G** | 4(14. 28%) | 6(60%) |  |
| **Allele** |  |  | <0.0001 |
| **C** | 44(78.57%) | 5(25%) |  |
| **G** | 12(21. 43%) | 15(27%) |  |

### 3.4 TP53 Nucleotide changes

DNA Sequence analysis was done by using Gene screen and BioEdit softwares, the following sequence variants were observed on comparison with the reference sequence (NG_017013). Mutations in the *p53* gene were found in 44%, 28% and 12% of esophageal squamous cell carcinomas, adenocarcinomas and small cell carcinomas, respectively. As shown in figure5, in exon 4, no mutation was detected other than NP_000537.3:p.R72P. While in exon 8, g.18783-18784AG>TT p.E285E, g.18803A>C p.K291T, g.18860A>C p.P301P, g.18845A>T p.K305M and g.18863_ 18864 InsT were observed. (Figure7)

**Figure 5.**
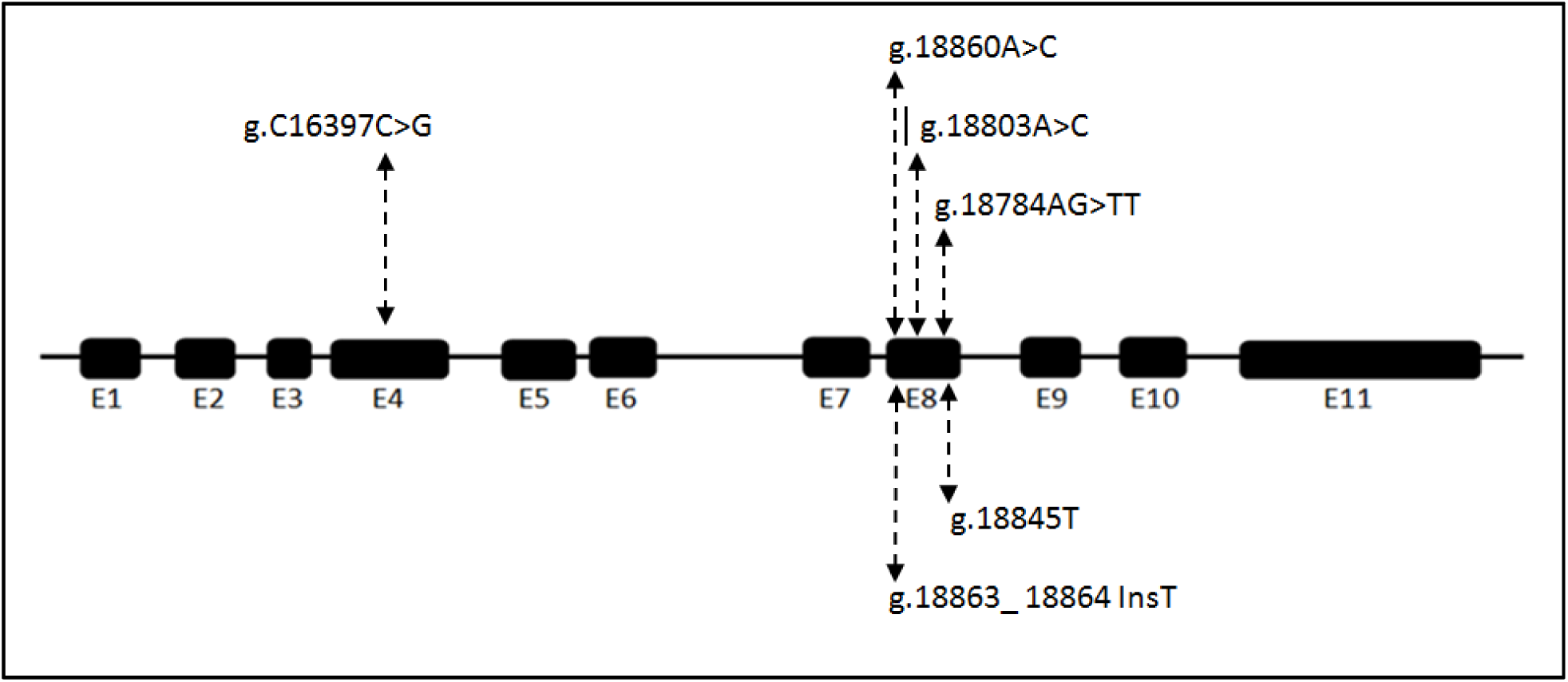
Illustrates the *TP53* gene and genetic alterations observed in exon4 and exon 8 for the studied population. E1 to E11 indicate exons 1 to 11.

Non-synonymous variants (R72P, K291T and K305M) were then functionally analyzed with SIFT, Polyphen-2, I-Mutant-3, and PhD-SNP to predicted their pathological effects, the results are provided in Table 4. The 3D structure of the variant K291T were obtained using Project Hope software, (Figure6c). While for R72P and K305M variants, we used Raptor X online software for prediction and Chimera for visualization, (Figure7c).

**Table 4.** Functional analysis of SNPs obtained by various sequencing software’s

| SNP | SIFT |  | Polyphen-2 |  | I-Mutant<br>prediction | RI | PhD-SNP |  |
| --- | --- | --- | --- | --- | --- | --- | --- | --- |
|  | Score | Prediction | Score | Prediction |  |  | Prediction | RI |
| **R72P** | 0.52 | Tolerated | 0.083 | Benign | Increase protein stability | 2 | Neural Polymorphism | 7 |
| **K291T** | 0.00 | deleterious | 0.972 | Probably damaging | Decrease protein stability | 4 | Neural Polymorphism | 0 |
| **K305M** | 0.00 | deleterious | 1.000 | Probably damaging | Decrease protein stability | 2 | Disease-related Polymorphism | 3 |

SNP: single nucleotide polymorphism; RI: reliability index

**Table 4.** Summary of published studies on Pro72Arg polymorphism in esophageal carcinoma in different populations

| Populations | Screening method | Inference | Reference |
| --- | --- | --- | --- |
| Sudanese | RFLP-PCR<br>DNA sequencing | ↑ risk with PP genotype | Present study |
| Sudanese | PCR | ↑ risk with RP genotype | (22) |
| South African | PCR-SSCP<br>DNA sequencing | ↑ risk with R allele | (49) |
| North Indian | PCR-RFLP | ↑ risk with RP genotype | (50) |
| Chinese | PCR-RFLP | ↑ risk with PP genotype | (51) |
| Chinese | PCR-SSCP<br>DNA sequencing | No association | (52) |
| **Chinese** | PCR-RFLP | ↑ risk with PP genotype | (53) |
| **Chinese** | TaqMan assay | ↑ risk with RR and RP genotype | (54) |
| **Caucasian** | TaqMan assay | ↑ risk with PP genotype | (55) |
| **American** | TaqMan assay | No association | (56) |
| **European and Asian** | Sequencing | ↑ risk with RR genotype | (57) |
| **Korean** | Real Time PCR | ↑ risk with P allele | (58) |
| **Japanese** | PCR | No association | (59) |
| **German** | PCR<br>DNA sequencing | ↑ risk with R allele and HPV<br>infection | (60) |
| **European** | APEX | No association | (61) |

**Figure 6:**
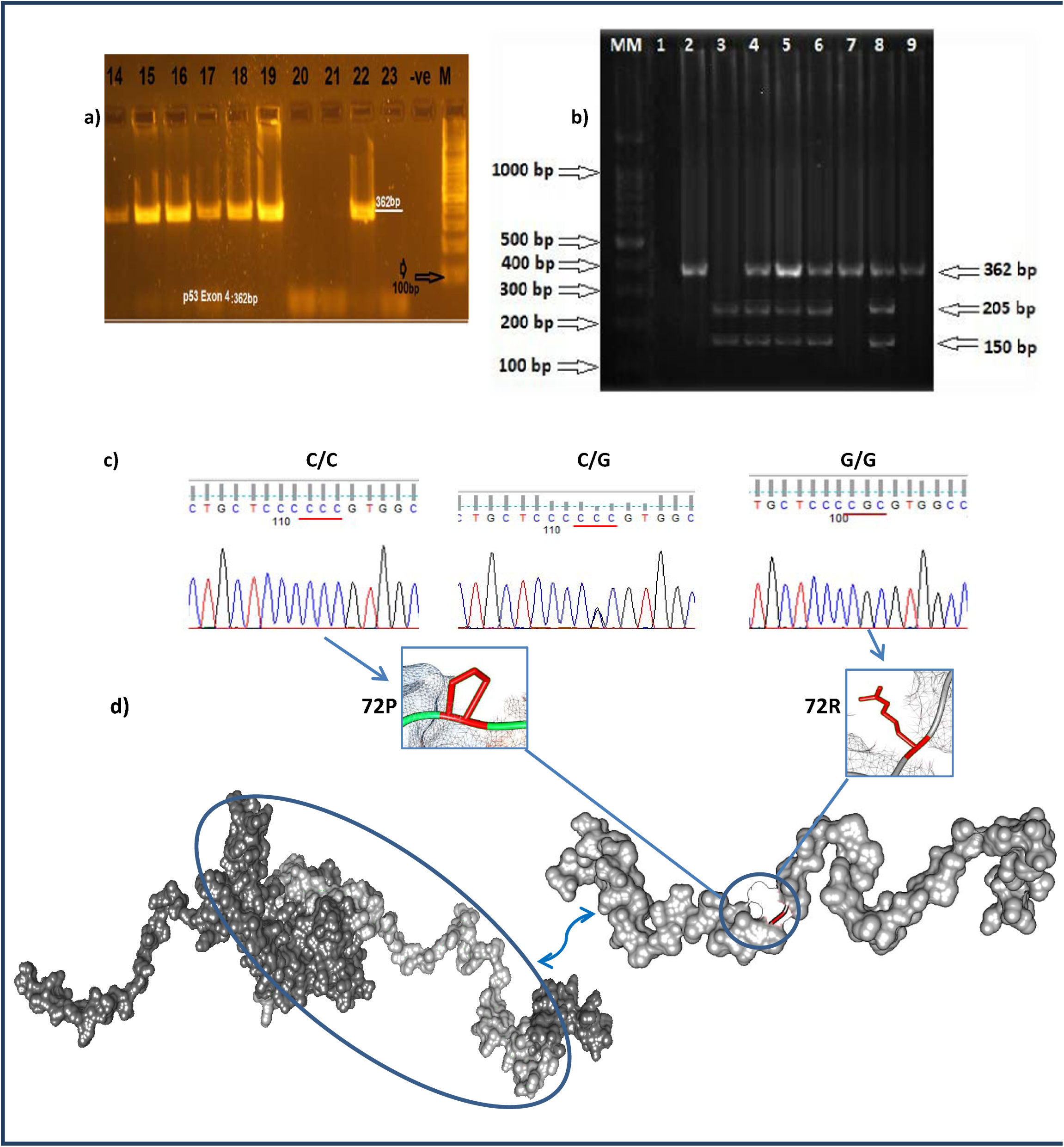
a) PCR amplification of p53 genes on 1.5% agarose gel electrophoresis. M DNA ladder: MW 100 bp lane 14-15-16-17-18-19-22showing typical band size of (362bp) corresponding to the molecular size of p53gene exon 4 and 20-21-23 none amplified. b)Genotyping of *TP53* gene for the codon 72 polymorphism by PCR-RFLP. The wild type G/G genotype produced two bands: 205bp and 150 bp. G/C genotype produced three bands: 362bp, 205bp and 250bp. C/C genotype produced single band 362bp. c) Sequencing chromatogram of codon72. d) 3D structure of TP53.

**Figure 7.**
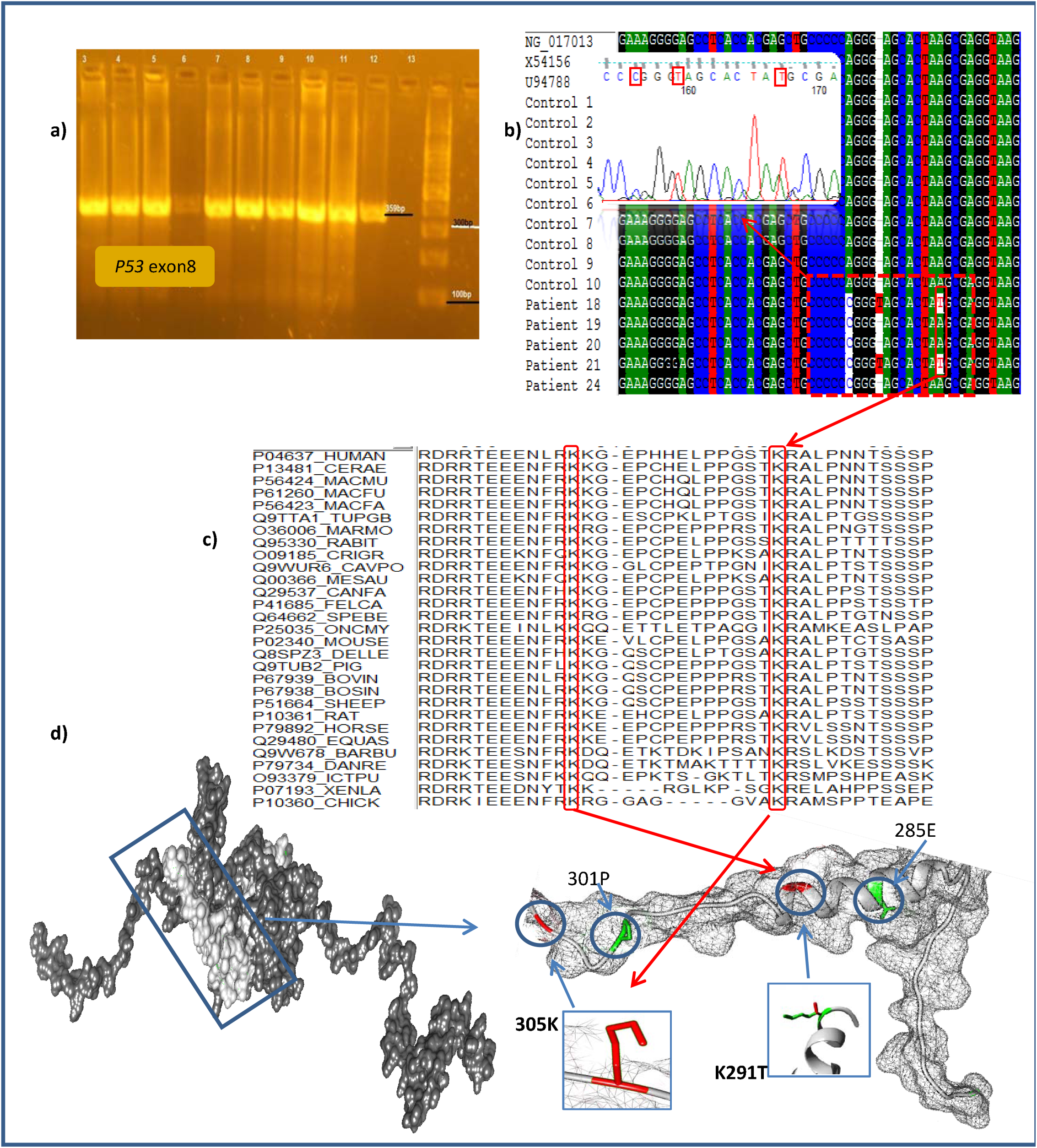
a) PCR amplification of *p53 gene* exon8 on 1.5% agarose gel electrophoresis. M DNA ladder: MW 100 bp. lanes 3-4-5-7-8-9-10-11-12 show typical band size of (359bp) corresponding to the molecular size of p53 gene exon8 and 9-13 none amplified. b) Shows the chromatogram of patients with small cell carcinoma. And BioEdit multiple sequences alignment determining mutations in exon 8. c) Multiple sequence alignment for p53 gene from different organisms. d) 3D structure of TP53.

## 4. Discussion

Despite different genetic and epigenetic alterations involving oncogenes activation, genetic variations in TP53 tumor suppressor represent the fundamental events related in both early and advanced stage of the esophageal tumor.(44, 45) Previously, TP53 mutations have been used as prognostic markers for patients’ response to treatment and/or outcome. (46, 47) In this study, we found a significant association between the accumulation of p53 protein and mutations in exon 4 and 8. Although, all patients with p53 mutations were immune-positive, there were five patients who were immune-positive but had no mutations in exon 4 and 8. Two of whom with adenocarcinoma. other study conducted by Doak *et al.* reported that the most immuno-positive adenocarcinoma cases had no demonstrable p53 mutation and immunohistochemistry is a poor indicator of p53 gene mutations.(48) Thus, further investigations are required to determine the underlying mechanisms that are responsible for the accumulation of the P53 protein.

In our study, we evaluated the frequency of the p53 Pro72Arg polymorphism in esophageal cancerous patients compared to healthy control subjects. Our findings exhibit a significant association between esophageal carcinoma and the Pro72 variant of the Pro72Arg polymorphism of the p53 gene. The Pro72 variant exhibits a higher level of G1 arrest and decreased apoptotic potential than the Arg72 variant.(13, 15) A number of studies have suggested that the Pro allele or the Arg allele of p53 codon 72 polymorphism had a significant effect on the risk of esophageal cancerogenesis while others did not demonstrate any significant association between them, as illustrated in Table 4.

The sequencing analysis of exon 4 for 28 patients, revealed no mutations or SNPs other than Arg72Pro. While in exon 8, we performed sequencing for 5 patients and we found silent mutation P301P shared in all of them. Further studies with large sample size are required to demonstrate its usefulness in the screening of EC. The known hotspot mutations in exon8 (p.C275Y, p.P278S and p. E298)(46) were not detected in this study.

The most important finding in this investigation was that two patients who diagnosed with small cell carcinoma have beside the previously mentioned silent mutation P301P, a novel insertion mutation (18736_18737 InsT) and missense mutation K305M. Small cell carcinoma of the esophagus (SCCE) is one of the deadliest aggressive cancers with poor prognosis.(62) It accounts for 1–2.8% of all esophageal carcinomas. Most diagnosed patients with SCCE die within 2 years and survival rates ranging between 8–13 months.(63) Histologically, SCCE is similar to SCC that arises in the lung and other extra-pulmonary organs. It is characterized by neuroendocrine-like architectural patterns, including nested and trabecular growth with common characteristics including peripheral palisading and rosette formation.(64) Understand the pathogenesis of SCCE is are urgently required to develop new diagnostic tools and effective treatment for this deadly cancer. Genetic alterations in exon 8 are shown in Table 5.

**Table 5.** Genetic alterations of exon 8 in esophageal carcinoma patients

| Patient No. | Type of carcinoma | Codon | Base change | Event | Mutation<br>Amino-acid substitution |
| --- | --- | --- | --- | --- | --- |
| **Patient 18** | Small cell carcinoma | 921 | AAG →ACG | Transition | Lys →Thr |
|  |  | 301 | CCA →CCC | Transition | Pro →Pro |
|  |  | 302_303 | GGG_Ins T_ | Frameshift* | Gly Ser →Gly Stop codon |
|  |  | 305 | AGC |  | Lys →Met |
|  |  |  | AAG →ATG | Transversion |  |
| **Patient 19** | Small cell carcinoma | 301 | CCA →CCC | Transition | Pro →Pro |
| **Patient 20** | Squamous cell carcinoma | 301 | CCA →CCC | Transition | Pro →Pro |
| **Patient 21** | Small cell carcinoma | 301 | CCA →CCC | Transition | Pro →Pro |
|  |  | 302_303 | GGG_Ins T_ | Frameshift* | Gly Ser →Gly Stop codon |
|  |  | 305 | AGC |  | Lys →Met |
|  |  |  | AAG →ATG | Transversion |  |
| **Patient 24** | Adenocarcinoma | 301 | CCA →CCC | Transition | Pro →Pro |

\* Novel mutation found in this study

The missense mutation K305M is located within a stretch of residues, bipartite nuclear localization signal, which is annotated as a special motif in UniProt (N6-acetyllysine). This mutation may disturb the motif and probably affect its function.(39, 65) Moreover, this mutation matches a previously described variant implicated in a familial cancer not matching LFS, germline mutation and somatic mutation.(39) A silent mutation at position 305 is also reported by Rihab *et al.* in Sudanese patients who were diagnosed with esophageal squamous cell carcinoma.(66)

In patient 18, we found missense mutation of a Lysine into a Threonine at position 291. This residue is located in a domain which is important for binding of other molecules and in contact with residues in a domain that is also important for binding (DNA binding site GO:0003677 and DNA-Binding Transcription Factor Activity GO:0003700). The mutation might disturb the interaction between these two domains and consequently affect the function of the TP53 protein.(39) Moreover, this mutation is located in a region with known splice variants, described in sporadic cancers and somatic mutation (dbSNP:rs372613518 and dbSNP:rs781490101) corresponds to variant. Additionally, mutagenesis experiments have been performed on this position and the next (291 and 292). Mutation of the wild-type residues (KK) into (RR) abolishes polyubiquitination by Makorin Ring Finger Protein 1 (MKRN1).(67) Also, patient 18 had a silent mutation at position 285. A germline mutation and somatic mutation in this position implicated with Li-Fraumeni syndrome (LFS) (OMIM:151623). Further studies with large sample size are required to demonstrate the usefulness of these mutations in the screening of EC especially SCCE. Studying the genetic alteration of esophageal carcinoma will help in the development of new diagnostic and therapeutic tools for its treatment.

## In conclusion

we found a significant association between the overexpression and an accumulation of TP53 protein; and mutation in exon 4 and 8. Also, there is a significant association between esophageal carcinoma and the Pro72 variant of the Pro72Arg polymorphism of the *p53* gene. A silent mutation *P301P* was detected in all of examined cases. Two patients who diagnosed with small cell sarcoma have shared the same mutations in exon8.

